# Understanding the physiological behaviour of disc cells in an *in vitro* imitation of the healthy and degenerated disc niche

**DOI:** 10.64898/2025.12.09.693181

**Authors:** Joseph Snuggs, Shaghayegh Basatvat, Exarchos Kanelis, Abbie Binch, Leonidas Alexopoulos, Marianna A. Tryfonidou, Christine Le Maitre

**Author notes:** Corresponding Author: Professor Christine Le Maitre, Division of Clinical Medicine, School of Medicine and Population Health, University of Sheffield, Sheffield, S10 2RX, UK.

## Abstract

**INTRODUCTION:** The natural environment within the nucleus pulposus (NP) is hypoxic, acidic, low in nutrition and exerts a high osmotic-pressure. NP cells have adapted to this harsh environment, however, *in vitro* conditions often fail to recapitulate this environment. Hence, this study aimed to develop media to mimic the conditions of the native NP, with regards to pH, osmolality, glucose, combined with culture in physioxia (5% O_2_), and determine their effects upon human NP cells and tissue.

**METHODS:** All media utilised low glucose DMEM/serum free conditions with supplements supporting matrix synthesis, based on media recommended for alginate culture (standard media). Healthy media modulated osmolarity (425mOsm/Kg), and pH7.2, degenerate media consisted of 325mOsm/kg and pH6.8. The latter was further supplemented with 100pg/ml IL-1β (degenerate+IL-1β media). NP cells in 3D alginate and NP tissue explants were cultured in these media for up to 2 weeks in physioxia (5% O_2_) to determine effects on viability, mitochondrial activity, protein expression and secretome.

**RESULTS:** Media osmolarity and pH remained stable and cell viability was not altered by any media composition. Mitochondrial activity was increased during short term cultures, whilst a decrease was seen following 14 days in degenerate media. The secretome of NP cells was differentially affected in healthy or degenerate media, with most increases in catabolic cytokines observed following the addition of IL-1β. Tissue explants showed stability of protein expression of matrix components in both healthy and degenerate+IL-1β media, with limited effects seen on the secretome.

**DISCUSSION:** The media formulations developed here can provide more appropriate environmental conditions *in vitro,* mimicking more closely the *in vivo* conditions observed within healthy and degenerate IVDs. The application of which can provide more appropriate culture conditions to test potential therapeutic approaches and understand more fully the pathogenesis of disease using *in vitro* and *ex vivo* models.

## Introduction

The adult IVD is the largest avascular organ with blood vessels located only within the annulus fibrosus (AF) and vertebral endplates^1^. The cells within rely on diffusion through the dense extracellular matrix (ECM) via load-induced water transport across the cartilaginous end plate (CEP) for the supply of nutrients (e.g. O_2_ and glucose) and removal of waste products (e.g. lactic acid)^2^. As a result, there are steep concentration gradients of oxygen, glucose and lactic acid in the IVD, with the nucleus pulposus (NP) being particularly low in glucose and oxygen (ranging from 1 - 5%^1,3,4^), whereas lactic acid levels are relatively high^4–7^. As such, the pH within the healthy NP is relatively low (pH7.2 – 6.9)^1,8^ and decreases further during IVD degeneration to ∼(pH6.8 – 6.5), whilst oxygen concentration remains low (∼5%)^8,9^. These changes are in part due to the calcification of the CEP, decreased nutrient diffusion, and compromised removal of waste products including lactic acid, which further lead to cellular dysfunction and decreased matrix synthesis^2,4,10^.

In the healthy IVD, the ECM of NP tissue is rich in glycosaminoglycans (GAGs). These negatively charged, hydrophilic proteoglycan sidechains enable water and cation retention within the NP, providing an environment that is highly hydrated and hyperosmotic. The NP is confined by the AF and the CEP, leading to increased osmotic and hydrostatic pressure enabling the cells residing with the disc to adapt to the biomechanical loading of the spine^11,12^ by increased production of ECM. However, during degeneration the GAG content of the NP is reduced, which leads to a decrease in tissue hydration and osmolality^13,14^. These changes contribute to IVD degeneration as cells can no longer adapt to their permanently altered osmotic environment^15^.

The function of NP cells is highly dependent on both oxygen concentration and pH, of note matrix production is increased when cells from the NP are cultured at 5% O_2_ ^16,17^, whilst reports of the effect of pH to mimic the native disc environment (pH7.2-6.9) have not been widely reported^1,18–23^. Whereby low pH on bovine NP cells decreased GAG synthesis but did not affect degrading enzyme production^18^, decreased viability and ECM production on porcine NP cells^23^, rat NP cells low pH promoted cellular senescence and decreased proliferation^19^, whilst on human NP cells it altered mechanoresponses^21^.

The observed changes to oxygen concentration, pH and osmolality during the degenerative process, could contribute to the transition from an anabolic to catabolic environment^24^. Cytokines and growth factors produced by NP cells^12,25–27^ play important roles in normal homeostasis and in the onset and progression of disc degeneration^12,28^. Importantly, locally produced IL-1β has been shown to drive degenerative features in human cells including decreased ECM synthesis and upregulation of matrix degrading enzymes, such as matrix metalloproteinases (MMP) and a disintegrin and metalloproteinase with thrombospondin motifs (ADAMTS)^29^. IL-1β has also been shown to act as a plethoric cytokine inducing the production of many other cytokines and chemokines by NP cells *in vitro* and thus can be deployed to induce the catabolic cytokine secretome seen during IVD degeneration^30^. Furthermore, IL-1β drives increased production of vascular endothelial growth factor (VEGF) and neurotrophic factor expression^31–33^, which are thought to contribute to the vascularisation and infiltration of sensory nerve fibres, potentially leading to sensitisation and pain^34–37^.

However, most *in vitro* environmental conditions to date have not recapitulated the combined environmental conditions of low nutrient, low pH, altered OSM, physiological concentrations of IL-1β and physioxic environment. Within the recently published consensus methodology for NP cell *in vitro* culture, standardization of extraction, expansion, and re-differentiation of NP cells was proposed with support across the spine community whereby recommendations were made for common species used for NP cell culture and research^38^. However, these recommended culture conditions can be further improved to mimic the environment of the NP better. Thus, to gain an improved understanding of cell behaviour and phenotype *in vivo*. In this study, we aimed to develop and compare *in vitro* culture conditions resembling the healthy and degenerated NP environment more closely by adjusting the pH, osmolarity, glucose concentration combined with culture under physioxia, furthermore addition of a physiologically relevant concentration of IL-1β was investigated^30,39^. The behaviour of NP cells and *ex vivo* tissue explants from human IVDs taken from microdiscectomy surgery were investigated within these novel culture conditions compared to standard alginate culture as previously recommended^38^.

## Methods

### Development of healthy and degenerated disc media

To mimic the IVD environment *in vitro*, this study adjusted the osmolality to match those reported for the non-degenerate and degenerate NP niche using the impermeant cation N-methyl-D-glucosamine HCL to prevent solute-specific target effects from other compounds^40^. To adjust the pH, the Henderson-Hasselbalch equation was used to determine the concentration of sodium bicarbonate required to accurately alter media pH when cultured in 5% CO_2_ ^21^. To mimic the low nutrition environment, low glucose (1g/L) DMEM was utilised. The media compositions used were based on those recommended in the recent consensus NP cell culture paper for 3D alginate culture which formed ‘standard’ media in this study^38^. Basal culture media for all compositions utilised the Fetal Bovine Serum (FBS) alternatives, exchanging FBS for Albumax, insulin-transferrin-selenium (ITS-X) and L-Proline^38^. Media was further supplemented with ascorbic acid to support collagen synthesis^38^. To prepare the media, powdered low glucose DMEM without pre-added sodium bicarbonate was utilised so that sodium bicarbonate concentration could be adjusted to alter the pH of the solution (Table 1). Degenerate media was further supplemented with a physiological concentration of recombinant human IL-1β (100pg/ml, Peprotech)^39^ to mimic the catabolic environment of the degenerate NP.

**Table 1:** Media composition.

| Product | Standard | Healthy | Degenerate | Supplier |
| --- | --- | --- | --- | --- |
| **Low glucose DMEM (1g/L Glucose + Pyruvate)** | 4.99 gram per 500 mL | 4.99 gram per 500 mL | 4.99 gram per 500 mL | Gibco:31600-083 |
| **Sodium bicarbonate** | 0.851 gram per 500 mL | 0.425 gram per 500 mL | 0.213 gram per 500 mL | Sigma: S-6014 |
| **N-Methyl-D-glucamine HCL (CAS 35564-84-4)** | 37.5 mM (exact amount to be | 92.5 mM (exact amount to be | 47.5 mM (exact amount to be | Santa Cruz: sc-286418 |

| **NaCl homologue (450mOsm/kg)** | determined with osmometer) | determined with osmometer) | determined with osmometer) |  |
| --- | --- | --- | --- | --- |
| **Pen/Strep** | 50 U/mL | 50 U/mL | 50 U/mL | Gibco:15070-063 |
| **Amphotericin B** | 2.5 µg/mL | 2.5 µg/mL | 2.5 µg/mL | Sigma: A2942 |
| **L-Ascorbic acid 2-phosphate sesquimagnesium salt hydrate** | 25 µg/mL | 25 µg/mL | 25 µg/mL | Sigma: A8960 |
| **L-glutamine** | 1% v/v | 1% v/v | 1% v/v | Gibco: 25030-024 |
| **ITS-X** | 1% v/v | 1% v/v | 1% v/v | Gibco: 51500-056 |
| **L-proline** | 40 µg/mL | 40 µg/mL | 40 µg/mL | Sigma: P5607 |
| **Albumax** | 1.25 mg/mL | 1.25 mg/mL | 1.25 mg/mL | Gibco: 11020-021 |

### Human NP samples

Human IVD samples were collected from patients undergoing microdiscectomy surgery with informed consent of the patients or relatives (Table 2). Ethical approval was granted from Sheffield Research Ethics Committee (IRAS 10226). Upon receipt, tissue was rinsed twice with 1x PBS containing 1% v/v penicillin/streptomycin (P/S) (Thermo Fisher, Warrington, UK). NP tissue was separated from AF/CEP and weighed in a sterile environment to enable wet weight normalisation. NP cell isolation and tissue digestion was performed as previously optimized and described in the consensus methodology^38^. Briefly, NP tissue was digested in type II collagenase (Thermo Fisher) at 64U/mL in lg DMEM (Thermo Fisher: 31600-083) containing 1% P/S and Amphotericin B (Sigma, Poole, UK) on a shaker (600rpm) for 4 hours at 37°C. Following digestion, the collagenase solution was passed through a 70µM cell strainer and centrifuged for 10 minutes at 400g. NP cells in monolayer were expanded up to passage two in complete medium containing hg-DMEM (HG-DMEM, Invitrogen: 10569010 HG) supplemented with 10% v/v heat-inactivated foetal calf serum (FCS, LifeTechnologies, Paisley, UK), 1% v/v P/S, 1% v/v L-Glutamine (Invitrogen: 25030-024), 2.5 µg/ml Amphotericin B (Sigma, A2942), 25µg/ml L-ascorbic acid 2-Phosphate (AA, Sigma, A8960) as described previously^38^, prior to encapsulation in alginate. To produce tissue explants, morphologically distinct NP tissue was dissected into 5 x 5mm diameter explants and cultured within semi-constrained silicone rings to limit tissue swelling during culture and retain their phenotype as demonstrated previously^41,42^.

**Table 2:** Patient details utilised for studies. M: Male; F: Female. Grade of Degeneration determined using NP score (0-9) from consensus human Grading scheme (DOI: doi.org/10.1002/jsp2.1167)

| Patient ID | Age | Sex | Disc Level | Grade of degeneration | Studies used for |
| --- | --- | --- | --- | --- | --- |
| HD596 | 21 | M | L4/5 | 5 | Viability assessment. |
| HD623 | 74 | M | L4/5 | 8 | Viability & Mitochondrial activity. |
| HD624 | 22 | F | L4/5 | 4 | Viability & Mitochondrial activity. |
| HD625 | 42 | M | L5/S1 | 6 | Viability & Mitochondrial activity. |
| HD585 | 34 | M | L4/5 | 5 | 3D alginate culture – Luminex analysis. |
| HD632 | 55 | M | L3/4 | 7 | 3D alginate culture – Luminex analysis. |
| HD639 | 32 | M | L4/5 | 6 | 3D alginate culture – Luminex analysis. |
| HD644 | 28 | F | L5/S1 | 6 | 3D alginate culture – Luminex analysis. |
| HD661 | 72 | M | L4/5 | 8 | 3D alginate culture – Luminex analysis. |
| HD741 | 41 | M | L4/5 | 3 | Tissue explant study. |
| HD742 | 66 | M | L4/5 | 4 | Tissue explant study. |
| HD744 | 32 | M | L5/S1 | 5 | Tissue explant study. |

### Encapsulation of Human NP tissue in alginate

Following expansion NP cells were encapsulated in 1.2% w/v medium viscosity alginate (Sigma: A2033) in 0.15M NaCl to a final cell density of 4 × 10^6^ cells/mL, and alginate beads formed. Alginate was polymerised in 200mM CaCl_2_ for 10 mins and washed in 0.15M NaCl as described previously^38^. NP cells were cultured in alginate beads for 2 weeks to regain their *in vivo* phenotype in 37°C 5% CO_2_, 5% O_2_ within standard alginate culture media (lg DMEM with Pyruvate (Thermo Fisher:31600-083), 50U/ml Pen/Strep (Thermo Fisher), 2.5 µg/mL Amphotericin B (Thermo Fisher), 25 µg/mL 2-AP-Ascorbic Acid (Sigma), 1% v/v L-Glutamine (Thermo Fisher), 1% v/v ITS-X (Thermo Fisher), 40 µg/mL L-Proline (Sigma) and 1.25mg/ml Albumax (Thermo Fisher))(Standard Media). Media was exchanged 3 times a week within a hypoxia glove box to maintain 5% O_2_.

### Media stability

Healthy and degenerate cell-free media was cultured at 5% O_2_, 5% CO_2_, 37°C for 7 days. At day 2, 5 and 7 media was removed and the pH was determined at 5% CO_2_ using a pH monitor within the hypoxia glove box, and osmolality was determined by freezing point depression using Model 3320 osmometer (Advanced Instruments, Dorset, UK).

### Cell viability testing

Following initial re-differentiation of NP cells within alginate beads for 2 weeks in standard media, beads were either maintained in standard media or transferred to healthy, degenerate or degenerate + 100pg/ml IL-1β media for 7 days. Metabolic activity of NP cells cultured within different media was assessed using the Alamar Blue (resazurin reduction assay) following 2 and 7 days in culture as previously published^43^. Furthermore following 7 days in culture, alginate beads were isolated, dissolved in Papain as described previously^44^ and DNA/alginate bead calculated using the Quant-iT™ PicoGreen™ dsDNA Assay Kit (P7589, Invitrogen), following the manufacturer’s protocol.

### Mitochondrial activity

Mitochondrial activity was assessed using the Image-iT™ Tetramethylrhodamine, methyl ester (TMRM) reagent (ThermoFisher). TMRM is a cell permanent dye which accumulates in active mitochondrial with intact membrane potentials. Whereby healthy cells with functioning mitochondrial result in a higher fluorescence reading, whilst upon loss of mitochondrial membrane potential, TMRM accumulation will halt, and signal intensity will decrease. TMRM is a dynamic dye that has a higher specificity for staining mitochondria compared to other available dyes^45^ and can be assessed by plate reader readout, avoiding imaging issues with 3D culture systems such as alginate. Furthermore, as there is no covalent attachment of TMRM to mitochondria it can move in and out of mitochondria dependant on changes in mitochondrial membrane potential making it ideal for dynamic analysis. In comparison to commonly utilised Mitotracker dyes which covalently attach to mitochondria and are retained even following subsequent drops in membrane potential, these are also more commonly used for microscopic imaging, which with 3D systems is problematic.

To determine that TMRM could indicate changes in mitochondrial membrane potential within human NP cells, alginate bead encapsulated NP cells (n = 3 patients: Table 2) cultured in standard media were treated with 10µM carbonyl cyanide-4-(trifluoromethoxy) phenylhydrazone (FCCP) for 10 mins prior to reading TMRM fluorescence compared to untreated controls. FCCP is a protonophore that disrupts the mitochondrial membrane potential and uncouples oxidative phosphorylation^45^. Following FCCP treatment the percentage of TMRM fluorescence (median value: 55.57%) was significantly reduced compared to untreated controls (median value: 99.5%) (*p*<0.0001) indicating a significant reduction in mitochondria membrane potential (Supplementary Figure 1).

To assess mitochondrial activity, human NP cells (n = 3 patients: Table 2) encapsulated in alginate beads were cultured with standard, healthy, degenerate or degenerate + IL-1β (100pg/mL) media at 5% O_2_, 5% CO_2_ for up to 2 weeks during re-differentiation. To determine the effects of media composition on mitochondrial activity at 1 and 2 weeks of re-differentiation cells were incubated with Image-iT™ TMRM Reagent (Thermo Fisher) at 0.1µM for 30 mins at 37°C. Alginate beads were then washed with PBS prior to reading fluorescence (ex: 548nm, em: 574nm) using a CLARIOstar plate reader (BMG Labtech, Aylesbury, UK). In follow up studies we also determined potential short-term effects of media composition on mitochondrial activity on NP cells for 10 and 30 mins following an initial 2-week re-differentiation period in standard media.

### Human NP cell secretome

Human NP cells isolated from 5 patient samples (Table 2) were cultured within alginate beads in standard media for 2 weeks to induce re-differentiation prior to maintenance within standard culture or transferred to healthy, degenerate, or degenerate media + 100pg/ml IL-1β for 72hrs to investigate their secretome into the culture media via Luminex assay. Conditioned media from alginate cultures was collected and analysed for 73 secreted proteins (Supplementary Table 1) using bead-based Luminex multiplex immunoassays (Protavio, Athens, Greece), utilising an eight-point standard curve as described previously^46^. Briefly, 96-well plates were coated with 50 µl of each 1X bead mix dilution (Mag-Plex® magnetic microspheres, Luminex Corp, Austin, TX, USA), containing 2,500 beads per bead ID, and incubated with 35 µL of standards, samples, and blanks for 90 mins at room temperature on an orbital shaker (1,000 rpm). Wells were washed twice with assay buffer (PR-ASSB-1x, Protavio, Greece), followed by the addition of 20 µl of a detection antibody mix at an average concentration of 1 µg/mL to each well, and incubated for 60 minutes at room temperature on an orbital shaker (1,000 rpm). After two more washes with assay buffer, 35 µL of Streptavidin-R-Phycoerythrin conjugate (5 µg/mL, SAPE-001, MOSS, USA) was added and incubated for 15 minutes under the same conditions. Finally, the wells were washed twice and resuspended in 130 µL of assay buffer, and the median fluorescence intensity (MFI) values were measured using the Luminex FLEXMAP 3D platform (Luminex Corp., Austin, TX, USA) with a minimum of 100 bead counts per sample. Protein concentration was normalised to patient-matched standard cultures to generate fold changes within healthy, degenerate and degenerate + IL-1β treated groups.

### NP explant protein expression and secretome

To determine influence of longer term exposure of cultures to healthy and degenerate+IL-1β media, and investigate with intact cell-ECM interaction. Human NP tissue explants were cultured immediately following removal from the body in semi constrained culture system for 14 days in standard, healthy or degenerate+IL-1β media prior to formalin fixation and paraffin embedding. The protein expression of collagen type I, collagen type II, aggrecan, IL-1β, IL-1R1, MMP3 and ADAMTS4 within NP explants was determined by immunohistochemistry on 4µm sections using previously published methods^47,48^. At least 200 cells per NP tissue explant were manually counted by 2 independent assessors blinded to group and determined as immunopositive (stained brown) or immunonegative (blue nuclear staining without the presence of brown cytoplasmic staining) and the percentage of cells positive for staining of the protein of interest was calculated. During the culture of NP explants, media was collected at 7-and 14-days. Media was changed every 3-4 days of culture and prior to collection at 7- and 14-days media was conditioned by NP explants for 3 days. The production of a panel of 18 secreted proteins (CCL2, CCL4, CX3CL1, CXCL1, CXCL10, GM-CSF, IL-1α, IL-1Ra, IL-6, IL-8, IL-17, MMP1, MMP3, MMP9, β-NGF, NT-3, TNF, VEGF-A) within cell culture supernatant was determined by Luminex Human Discovery Assay (18-Plex, LXSAHM-18, R&D Systems) using a Luminex™ 200™ Instrument System (Invitrogen) following the manufacturers protocols. Luminex data was analysed using the ProcartaPlex Analysis App (Thermo Fisher).

### Statistical analysis

All data were analysed using GraphPad Prism v10.5.0 (Dotmatics). Normality testing determined that media conditions (pH and OSM) and metabolic activity were normally distributed whilst DNA content, mitochondrial activity, Luminex and immunohistochemical analysis were non-normally distributed. Statistical differences between media conditions (pH and OSM) across time was investigated using a 2-way ANOVA with Tukey multiple comparison test, whilst metabolic activity (resazurin assay) was analysed with a 1-way ANOVA and Dunnett’s multiple comparisons test. Kruskal-Wallis with Dunn’s multiple comparisons test was used to determine influence of media composition on mitochondrial activity and mitochondrial activity across time in each media composition. Volcano plots for differentially secreted proteins were generated using GraphPad Prism v10.5.0 as data failed to follow a non-parametric distribution, Multiple Mann–Whitney U tests with a false discovery rate (FDR) using Two-stage step-up (Benjamini, Krieger, and Yekutieli) and FDR set to 5% was employed to identify statistically significant differences between the conditions depicted in each volcano plot. Volcano plots indicated significant differences between treatments (*p*<0.05) whilst potential trends were indicated where (*p*>0.05<0.1). Luminex analysis was plotted as actual concentrations produced but data was normalised to patients own standard media cultures for statistical analysis to determine fold changes from standard media and account for patient variability in baseline expression. Percentage immunopositivity was analysed with a Kruskal-Wallis with Dunn’s multiple comparisons test to determine significant differences across 3 or more groups. *p* ≤ 0.05 were considered as statistically significant, whilst trends were considered whereby *p*>0.05<0.1. Adjusted *p* values were used in all cases.

## Results

### Media conditions remain stable during culture

The stability of healthy and degenerate media over a 7-day period in culture without exchange of media was first investigated, determining pH and osmolality at 37°C, 5% O_2_, 5% CO_2_ demonstrating pH (Figure 1a) and osmolality (Figure 1b) remained stable for the first 5 days in culture with no significant difference in pH or OSM, whereafter levels significantly increased (pH: *p*=0.0039; osmolarity *p*=0.0024), demonstrating culture media should be exchanged prior to this time course. As culture media is exchanged every 2-3 days this was considered stable for culture.

**Figure 1.**
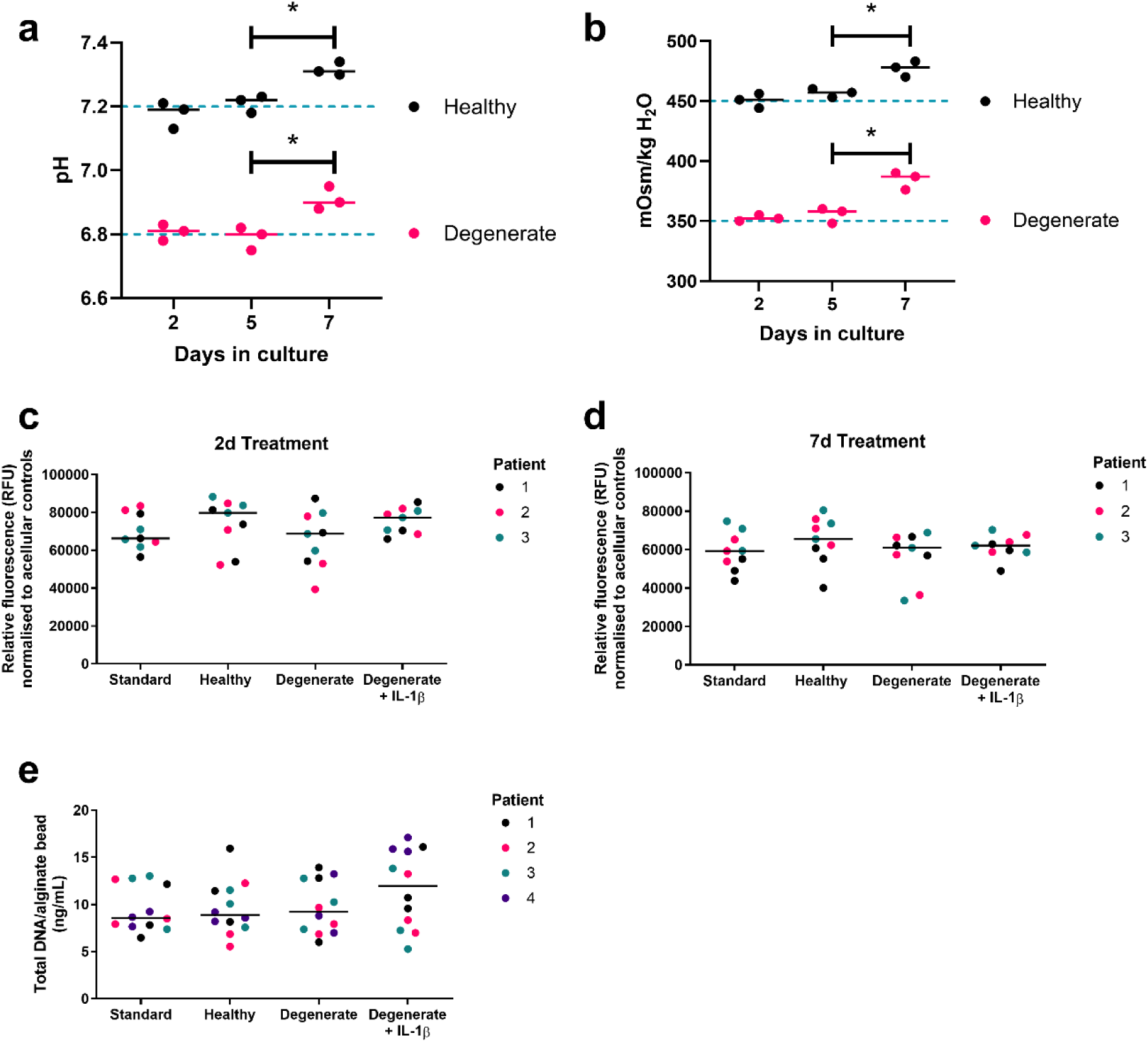
Media stability and viability of 3D alginate bead cultured human NP cells in response to treatment media. Change in (a) pH and (b) osmolality of healthy and degenerate media over 7d in culture at 5% O_2_ (37°C, 5% CO_2_). Metabolic activity of human NP cells following (c) 2d, and (d) 7 days treatment with standard, healthy, degenerate and degenerate + 100pg/mL IL-1β. Prior to treatment, NP cells were cultured in standard media for 2 weeks following suspension into alginate beads, to allow the 3D re-differentiation of cells. (e) Following 2 weeks of 3D re-differentiation and 7d treatment, alginate beads were dissolved in papain and total DNA/alginate bead was determined using Quant-iT™ PicoGreen™ dsDNA Assay Kit (P7589, Invitrogen), following the manufacturer’s protocol. *=*p*≤0.05.

### Human NP viability is retained during culture

The metabolic activity of human NP cells encapsulated in alginate beads and cultured within healthy or degenerate media with or without supplementation of 100pg/ml IL-1 was not adversely affected compared to standard media following 2d (Figure 1c) and 7d (Figure 1d) of culture with no significant difference between media conditions at either time point (*p*>0.05). Following 2w re-differentiation in alginate beads and a subsequent 7d treatment with standard, healthy, degenerate or degenerate + IL-1β media total double-stranded DNA content per alginate bead remained unaltered across all treatment groups (Figure 1e)( *p*>0.05), indicating that viability was maintained within all culture media tested.

### Mitochondrial activity

Mitochondrial activity of NP cells was significantly increased following 10 mins of short-term treatment with healthy and degenerate + IL-1β media following an initial 2 week re-differentiation period in standard media, compared to standard media controls (*p*<0.05)(Figure 2a). This effect was transient and after 30 mins of treatment no significant differences in mitochondrial activity were observed (Figure 2b). Whilst in cultures whereby re-differentiation was performed within standard, healthy or degenerate media, following 1 week re-differentiation in 5% O_2_ mitochondrial activity was significantly increased in healthy (*p*=0.027) and degenerate (*p*<0.0001) media compared to standard media controls and significantly decreased in degenerate + IL-1β (*p*=0.027) compared to degenerate media (Figure 2c). After 2 weeks of re-differentiation in 5% O_2_ mitochondrial activity was significantly decreased in degenerate media compared to healthy media (*p*=0.0003)(Figure 2d). During re-differentiation within standard, healthy or degenerate media+IL-1β in 5% O_2_ there was no significant difference in mitochondrial activity between 1 and 2 weeks (*p*<0.05), however a significant decrease was observed in degenerate media from 1 week to 2 weeks in culture (*p*<0.0001)(Figure 2c&d).

**Figure 2.**
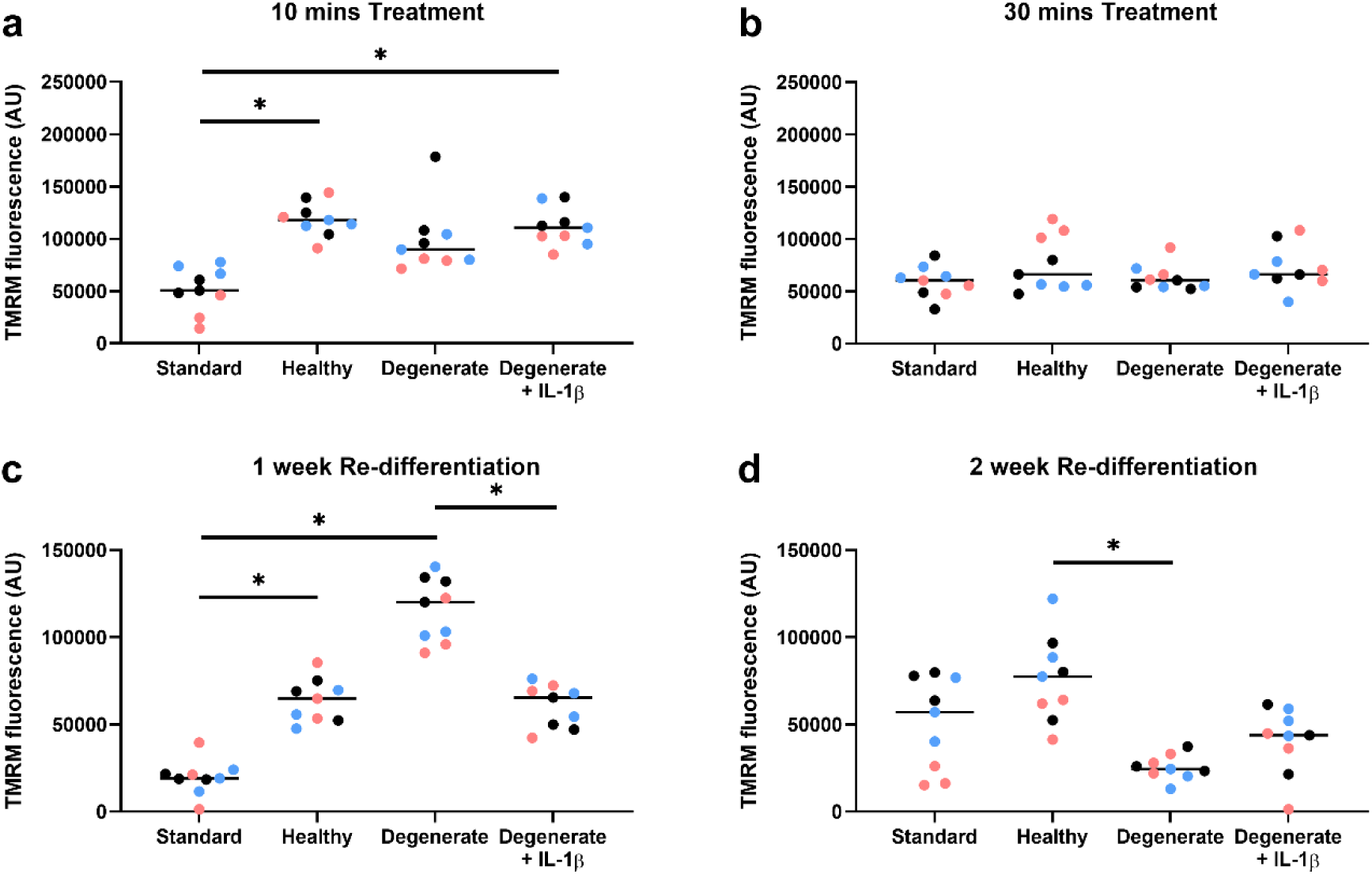
Mitochondrial activity. Mitochondrial activity determined using Image-iT™ TMRM Reagent in alginate bead-encapsulated human NP cells following short term (10, 30 mins) and long term (1, 2 weeks) treatment with standard, healthy, degenerate and degenerate + 100pg/mL IL-1β media cultured in 5% O_2,_ 37°C, 5% CO_2_ for the duration of culture. Individual patient data points shown in differential colours. *=*p*≤0.05.

### Secretome of Human NP cells cultured in alginate constructs in differential media

Luminex analysis of conditioned media from human NP cells extracted from degenerate discs, expanded and redifferentiated in alginate culture detected 55 secreted proteins out of the panel of 73 proteins investigated (Supplementary Table 1). Culture of human NP cells in media mimicking the healthy disc environment for 72hrs showed significant decreases in NGF, VEGF, PAL1, PROK1 and IL-10 compared to standard culture media (*p*<0.05)(Figure 3 + Supplementary Figure 2), with a trend for a decrease in IL-1α (*p*=0.066)(Figure 3). A significant increase was seen for CCL-7, M-CSF, CXCL11 in cultures with healthy media for 72 hrs compared to standard media (*p*<0.05)(Figure 3 + Supplementary Figure 2) and a trend for an increase in CCL20 (*p*=0.093) and CTACK (*p*=0.093). Fewer significant differences were observed when NP cells cultured in alginate were exposed to degenerate media for 72hrs compared to standard media with significant decreases seen for NGF, VEGF, PROK1, and MMP 2 (*p*<0.05)(Figure 3 + Supplementary Figure 2) with a trend for decrease in IL-6 (*p*=0.055) and CX3CL1 (*p*=0.1)(Figure 3) with only M-CSF significantly increased by culture in degenerate media compared to standard media (*p*<0.05)(Figure 3 + Supplementary Figure 2). Whilst 11 proteins were significantly increased within cultures with degenerate media + 100pg/ml IL-1β compared to degenerate media alone, namely: IL-1β, PROK1, NGF, IL-6, IL-11, IP-10, IL-20, M-CSF, CCL3, LIF and IL-1α (*p*<0.05)(Figure 3 + Supplementary Figure 1) with a trend seen for CXCL11 (*p*=0.065)(Figure 3), there were no proteins significantly decreased (Figure 3 + Supplementary Figure 2).

**Figure 3:**
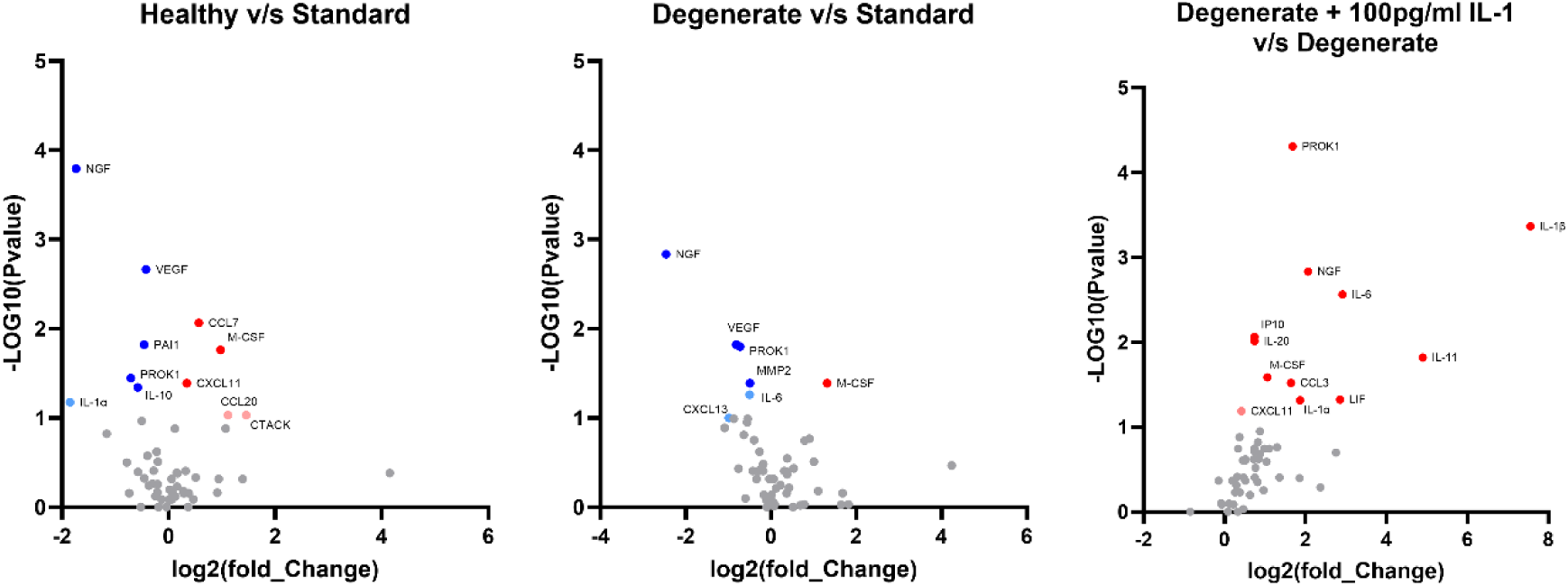
Secretome release from 3D cultured NP cells. Volcano plots indicating secretome of human NP cells cultured in alginate, following 2 weeks re-differentiation in standard culture media for 2 weeks at 5% O_2,_ 37°C, 5% CO_2,_ prior to transfer to standard, healthy, degenerate or degenerate + 100pg/ml IL-1β media for a further 72hrs. 73 Secreted proteins analysed with 55 secreted proteins detected within quantifiable levels. Volcano plots showing Log_2_(Fold change) against -Log_10_(P) within: A: NP cells cultured in healthy media compared to standard media; B: NP cells cultured in degenerate media compared to standard media; C: NP cells cultured in degenerate + 100pg/ml IL-1β compared to degenerate media alone. Dark red dots indicate significant increased proteins (*p*<0.05), and dark blue significant decreased proteins (*p*<0.05), pale red and blue indicate proteins which showed a trend for increase or decrease proteins (*p*<0.1 but >0.05).

### NP explant protein expression and secretome

The native expression of collagen type I, collagen type II, aggrecan, IL-1β, IL-1R1, MMP3 and ADAMTS4 within human NP tissue explants was not significantly different following 2 weeks of culture within healthy or degenerate + IL-1β media compared to standard media (Figure 4&5). However, in 2 of the 3 patient cultures (HD741 & HD742-Table 2) a significant increase was observed in the percentage of cells immunopositive for IL-1β and ADAMTs4 in explants cultured in degenerate + IL-1β media compared to standard media (IL-1: *p*=0.036; ADAMTs4: *p*=0.049) (Figure 5). Fifteen out of the 18 secreted proteins investigated were detectable (IL-1α, IL-1Ra, IL-6, IL-8, TNF, CCL2, CCL4, CXCL1, CXCL10, GM-CSF, MMP1, MMP3, MMP9, NT-3, NGF, VEGF)(Figures 6-9) and analysed with the aid of volcano plots to determine significant differences and potentially biologically relevant fold changes between culture media compositions (Figure 6). Following 7 days of culture only NGF was significantly decreased in healthy compared to standard media (*p*=0.02)(Figure 6&9), however this effect was lost following 14 days in culture. In NP tissue explants cultured in degenerate media supplemented with 100pg/ml IL-1β, NGF was also significantly decreased compared to standard media following 7 days and 14 days (7D:*p*=0.022; 14D:*p*=0.024)(Figure 6&9). CCL4 secretion was also significantly decreased (*p*=0.047)(Figure 6&8), whilst CXCL10 was significantly increased following 7 days culture of NP explants in degenerate + IL-1β media compared to standard media (*p*<0.047)(Figure 6&8). By 14 days of culture, CCL4 and CXCL10 did not differ between culture conditions. IL-8 was significantly increased in both healthy and degenerate + IL-1β media compared to standard media following 14 days in culture (Healthy: *p*=0.019; Degenerate+ IL-1β:*p*=0.01)(Figure 6&7).

**Figure 4.**
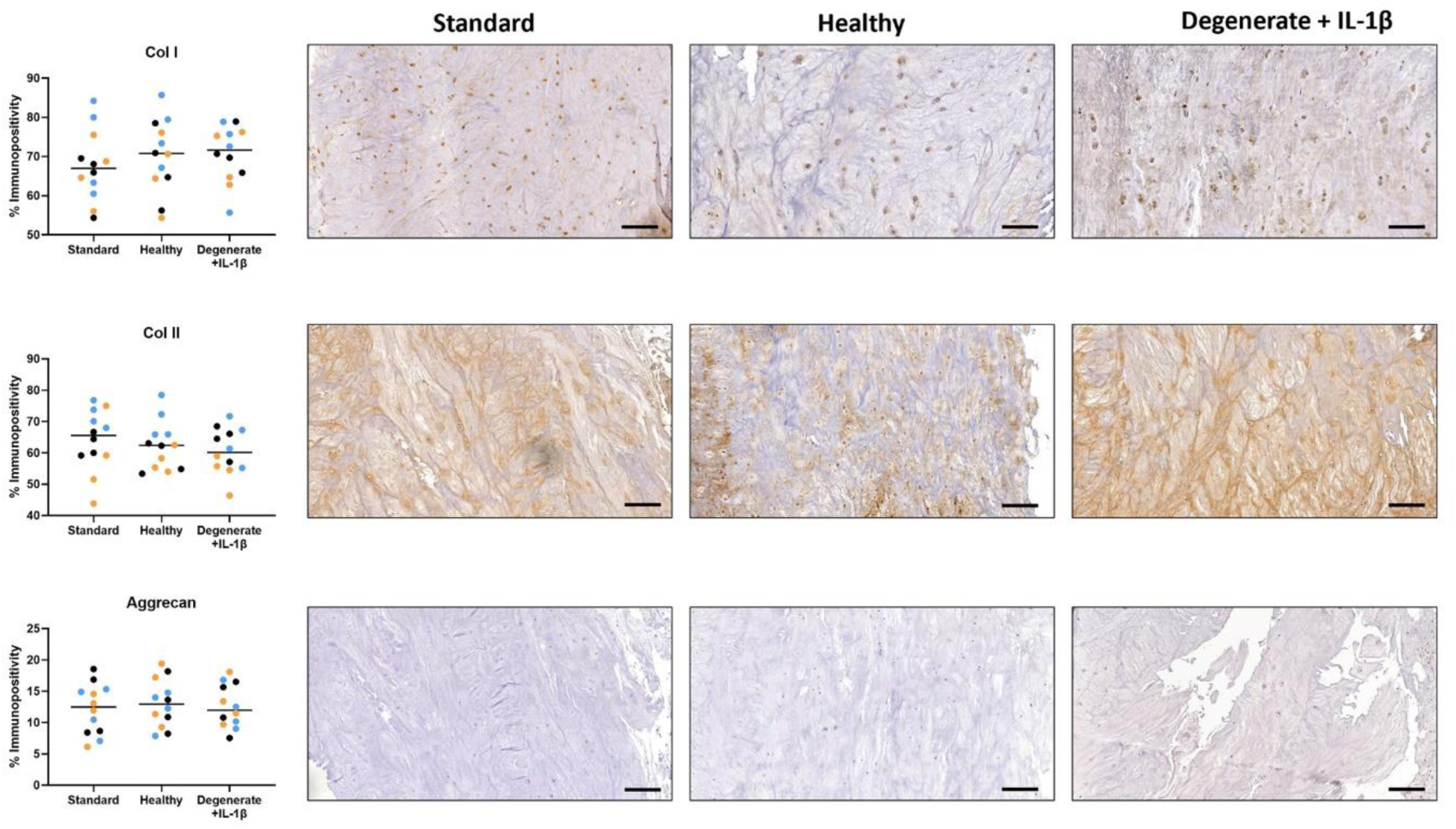
NP explant matrix protein expression. Expression of Collagen type I, collagen type II and aggrecan within *ex vivo* human NP tissue explants, determined by IHC, following 2 weeks of culture in standard, healthy, and degenerate + 100pg/mL IL-1β media at 5% O_2,_ 37°C, 5% CO_2_ for the duration of culture. Scale bar 100µm. *=*p*≤0.05.

**Figure 5.**
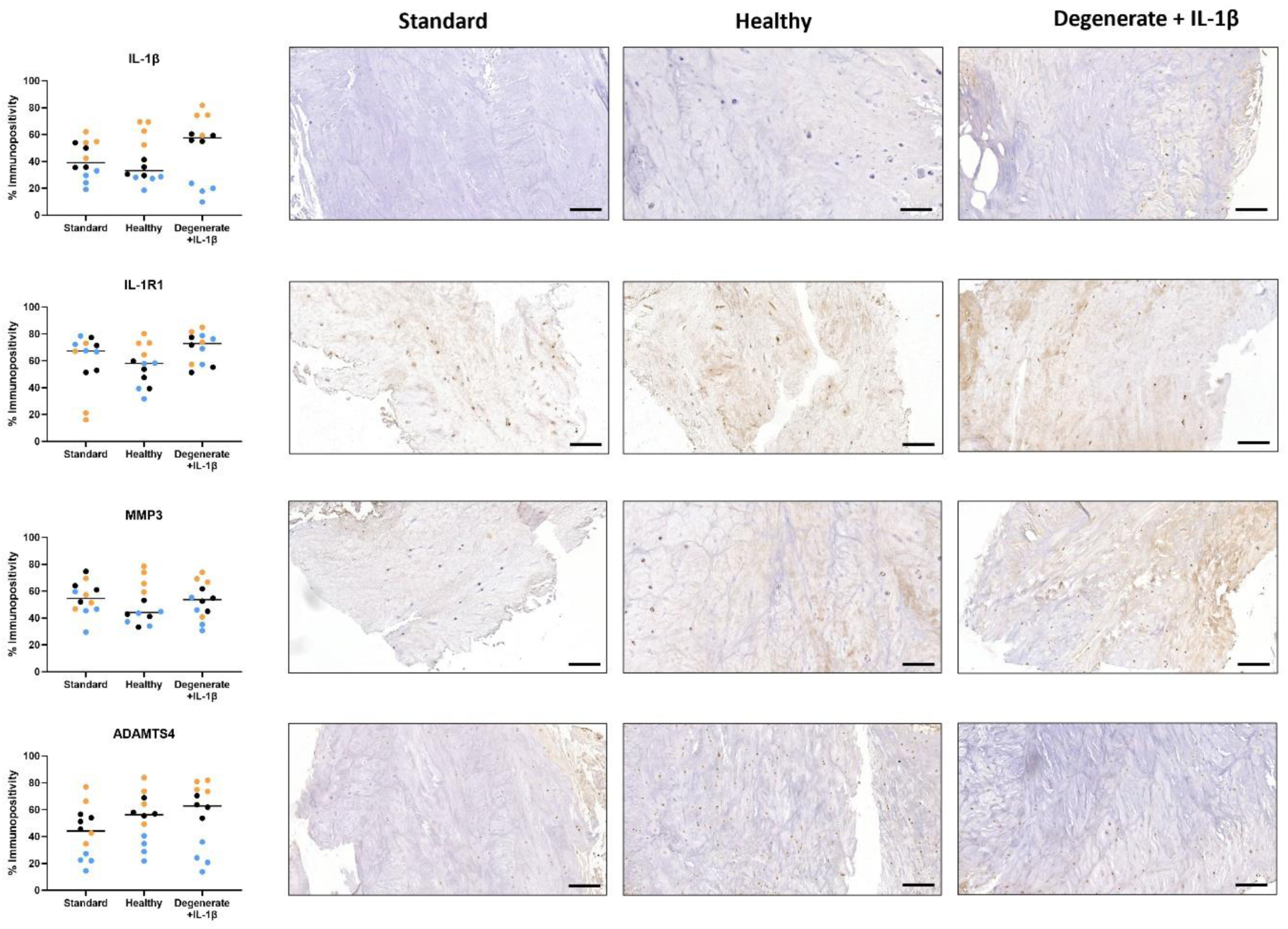
NP explant catabolic protein expression. Expression of IL-1β, IL-1R1, MMP3 and ADAMTS4 within *ex vivo* human NP tissue explants, determined by IHC, following 2 weeks of culture in standard, healthy, and degenerate + 100pg/mL IL-1β media at 5% O_2,_ 37°C, 5% CO_2_ for the duration of culture. Scale bar 100µm. *=*p*≤0.05.

**Figure 6:**
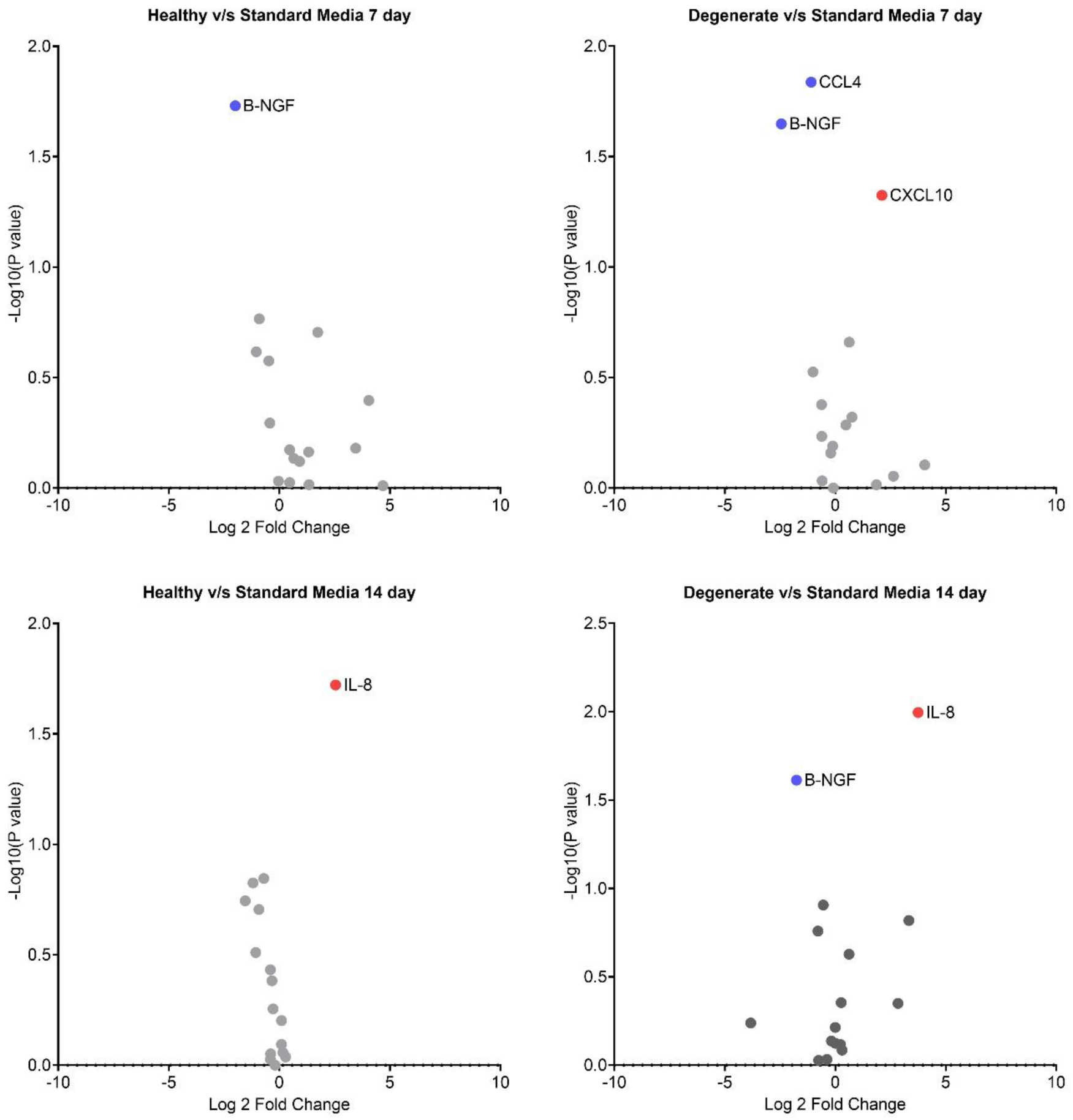
Secretome of NP tissue explants. Volcano plots indicating secretome of human NP tissue culture, immediately followed extraction, tissue explants cultured in semi-constrained silicone constructs for 2 weeks in either standard, healthy or degenerate + IL-1β culture media, 5% O_2,_ 37°C, 5% CO_2_. Eighteen selected secreted proteins were analysed with Luminex, with 15 proteins detected within quantifiable levels. Volcano plots showing Log_2_(Fold change) against -Log_10_(*p*) within: A: NP cells cultured in healthy media compared to standard media following 7days; B: NP cells cultured in degenerate + IL-1β media compared to standard media following 7 days; C: NP cells cultured in healthy media compared to standard media following 14days; D: NP cells cultured in degenerate + IL-1β media compared to standard media following 14 days; Dark red dots indicate significant increased proteins (*p*<0.05), and dark blue significant decreased proteins (*p*<0.05).

**Figure 7:**
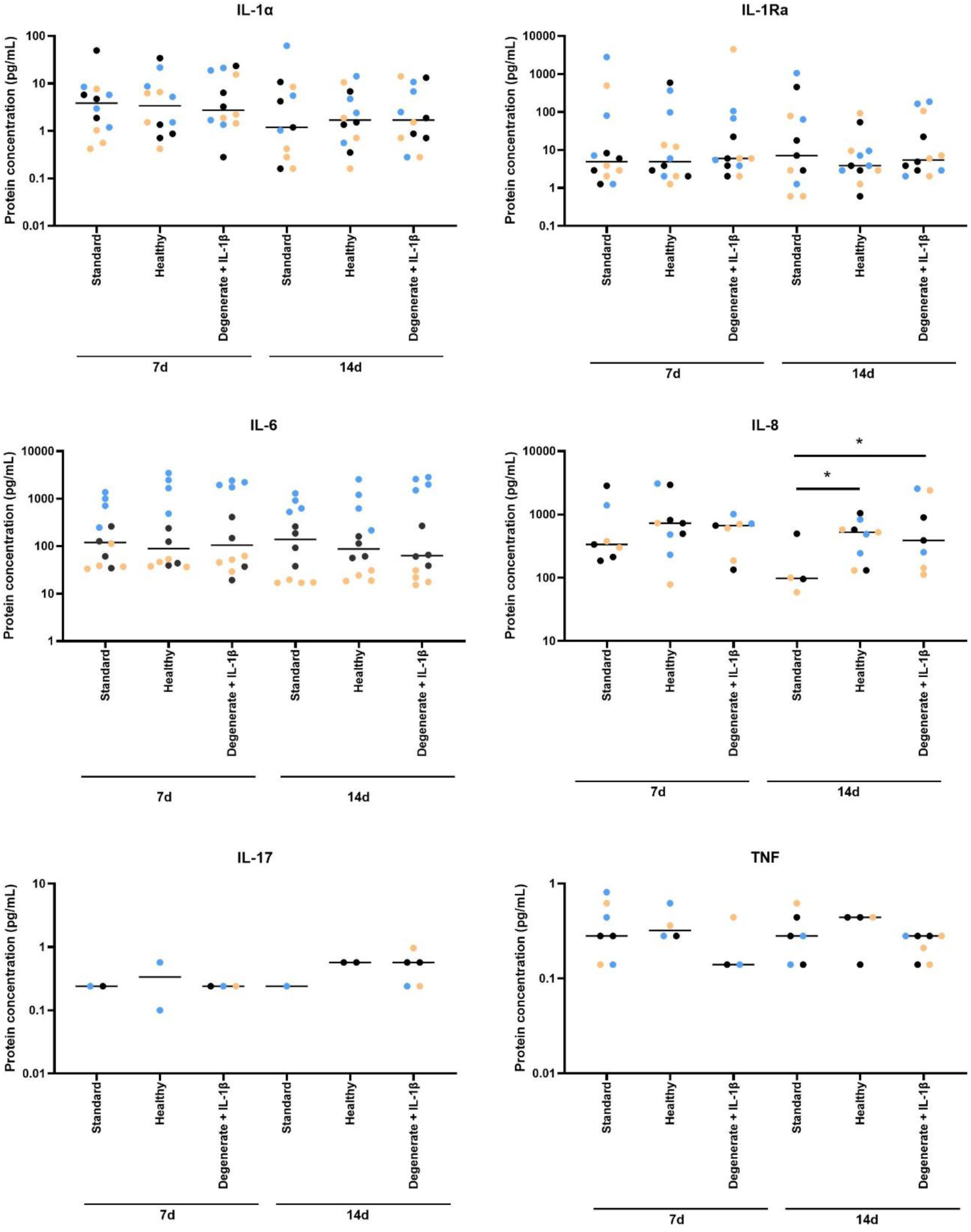
NP explant cytokine production. Production of cytokines (IL-1α, IL-1Ra, IL-6, IL_8, IL-17 and TNF) by *ex vivo* NP explants, using Luminex, following 7d and 14d culture within standard, healthy, and degenerate + 100pg/mL IL-1β, 5% O_2,_ 37°C, 5% CO_2,_ *=*p*≤0.05.

**Figure 8:**
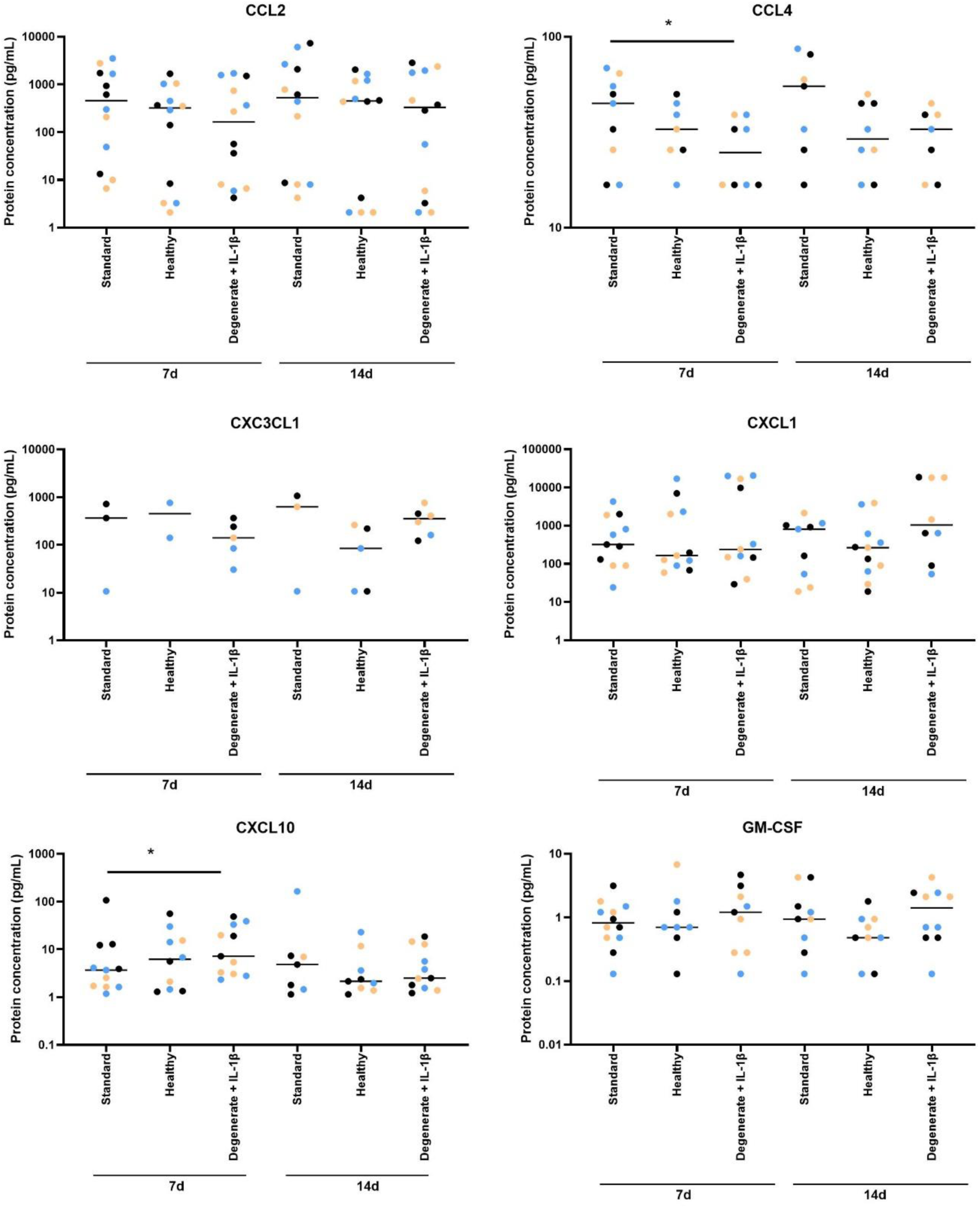
NP explant chemokine production. Production of chemokines (CCL2, CCL4, CX3CL1, CXCL1, CXCL10, GM-CSF) by *ex vivo* NP explants, using Luminex, following 7d and 14d culture within standard, healthy, and degenerate + 100pg/mL IL-1β, 5% O_2,_ 37°C, 5% CO_2,_ *=*p*≤0.05.

**Figure 9:**
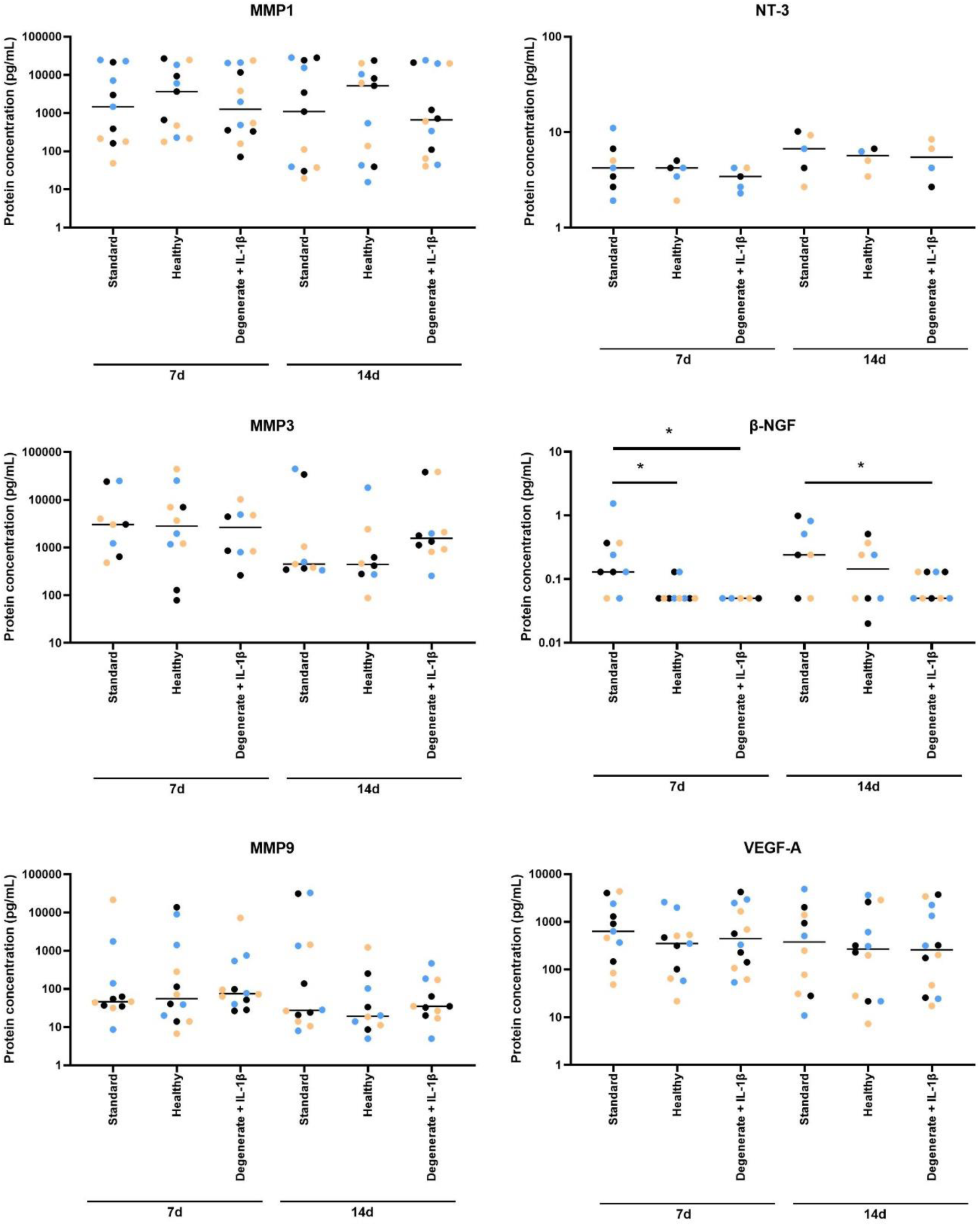
NP explant matrix degrading enzyme and neurotrophic/angiogenic factor production. Production of matrix degrading enzymes (MMP1, MMP3 and MMP 9), Neurotrophic factors (NT-3 and β-NGF) and vascular factor (VEGF) by *ex vivo* NP explants, using Luminex, following 7d and 14d culture within standard, healthy, and degenerate + 100pg/mL IL-1β, 5% O_2,_ 37°C, 5% CO_2,_ *=*p*≤0.05.

## Discussion

Here, media compositions to mimic the healthy and degenerate disc have been developed, whereby pH and osmolarity were shown to remain stable for at least 5 days without media exchange. Human NP cells cultured within 3D alginate cultures and NP tissue explants were cultured without any detectable loss of metabolic activity and DNA content suggesting retention of viability. Upon redifferentiation in standard media and changing into the degenerate media mitochondrial activity increased temporarily to decrease thereafter compared to healthy media culture. A comprehensive analysis was firstly performed on a broad secretome panel to assess the influence of such media over a short-term culture (72hr) on human NP cells previously expanded in hgDMEM and redifferentiated in standard lgDMEM. Whereby the secretome of NP cells was differentially affected in healthy and degenerate media, most notably decreased production of angiogenic and neurotrophic factors were seen compared to standard media. Whilst addition of physiological levels of IL-1 induced some of these factors alongside cytokines and catabolic factors. This was followed by investigating the influence of such media on the protein expression and secretome of NP tissue explants, whereby they were cultured within these physiological conditions immediately following isolation from the body. Within explants secretome effects were more limited and appeared to be donor dependant.

NP cells have shown limited response to changes in oxygen concentration or glucose conditions, within the physiological range^17^, and NP viability in prior studies where altered pH conditions that resemble mild to severe degenerative IVD, has been sustained^20^. Together with the results of the current study this suggests that as this is the native environment for NP cells they are primed to adapt and survive in these conditions. We previously showed that in human NP cells cultured in 3D alginate cultures, phenotype was maintained when cultured in low glucose, serum free conditions, which formed the basis of the standard culture media used in the current study^38^. Within the current study mitochondrial activity of NP cells in 3D alginate culture was initially increased in healthy and degenerate media both following short term treatment (10 mins) and during re-differentiation following 7 days. While this increased mitochondrial activity was maintained at 2 weeks of culture in healthy media, it was temporary in degenerate media, reducing well below the levels of healthy media following 14 days. Mitochondrial activity of human AF cells has been shown to decrease during IVD degeneration ^49^ and whilst the influence of culture conditions on human NP cell mitochondrial activity has not been investigated previously, mitochondrial activity has been shown to be altered in response to stimuli such as alterations in glucose concentration^50^, osmotic stress^51^, hydrostatic pressure^52^, pH and ion exchange^53^ within multiple cell types indicating mitochondrial activity changes observed here may be an adaptive response to specific alterations in media composition and may also be finely-tuned in response to environmental cues within the IVD. The findings here may implicate the degenerate environment in the decrease in mitochondrial activity seen during degeneration.

Despite the induction of metabolic activity observed in the 1^st^ week of culture, re-differentiated human NP cells cultured in 3D alginate beads in both healthy and degenerate media for 72hrs secreted lower levels of angiogenic and nerve ingrowth/sensitisation factors (NGF, VEGF, PROK1) compared to standard media. Given that OSM is similar between standard and degenerate media, but pH is decreased in both healthy and degenerate media compared to standard this suggests that this effect may be pH mediated. However following addition of physiological levels of IL-1β to degenerate media secretion of NGF and PROK1 increased, matching more closely the increased expression of neurotrophic factors seen in the degenerate disc^31^.

IL-1β has been shown in numerous studies, to induce the expression of cytokines, chemokines, matrix degrading enzymes and neurotrophic and angiogenic factors^12^, however most of these studies have utilised high glucose media often under 21 % O_2_ and many at higher concentrations^12^. In contrast this study aimed to mimic the natural disc environment by using low glucose conditions and physioxia (5%), where a number of factors were significantly increased (namely: IL-1β, PROK1, NGF, IL-6, IL-11, IP-10, IL-20, M-CSF, CCL3, LIF and IL-1α). However, for many factors responses were dampened or lost compared to previous studies. Within the current study IL-1β was deployed at a concentration aimed to mimic the concentration produced naturally by the degenerate disc cells. These concentrations were proposed from prior secretome studies showing that degenerate disc explant cultures attain 100pg/ml in culture without prior stimulation^39^. Such concentrations have been shown previously to induce cytokine and chemokine secretion in high glucose, high oxygen environments within alginate cultures^30^. As such the lack of responses for many factors such as the MMPs investigated here, which has been reported previously even at low concentrations of IL-1β^30^ demonstrates differential responses in low glucose, low oxygen, altered OSM and decreased pH conditions. It is possible that the metabolic activity of the disc cells observed within these conditions are lower than the abnormal environment induced by high glucose and high oxygen conditions, however to the authors knowledge this has not been determined to date. Furthermore, within cultures with high glucose and high oxygen it is probable that these environmental conditions would increase the concentration of advanced glycation end products due to glucose concentrations which are higher than that seen during diabetes^54^, with associated mitochondrial dysfunction^55^. Even more so, high oxygen concentrations will induce increased mitochondrial derived oxygen free radicals^56^ promoting catabolism. As such previous studies investigating the influence of cytokines and matrix degrading enzymes and other factors could be confounded by a high stress state of the cells^12^ cultured in the commonly used high glucose media and thus may amplify potential effects, highlighting the importance of evaluating potential pathogenic factors *in vitro* in an environment which is more representative of the *in vivo* environment.

The secretome of NP cells maintained in the native disc tissue environment was not differentially affected by healthy or degenerate+IL-1β culture conditions. There were however distinct donor differences, with increases only observed in number of immunopositive cells for IL-1β and the aggrecanase: ADAMTS4 in 2 of the 3 patients, the reasons for patient variability are difficult to discern given the low donor number, although the donor which failed to respond was younger but also slightly higher grade of degeneration than the responding donors. Although increased secretion of CXCL10 and IL-8 but decreased secretion of CCL4 and NGF was observed. This reduction in NGF production was evidenced in tissue explants cultured in healthy or degenerate+IL-1β media for 7 days and in degenerate+IL-1β media for 14 days, as well as, in 3D cell cultures in healthy and degenerate media for 72hrs. These findings seem in contradiction to disc degeneration where NGF is increased^31,33^ and has been shown to be induced by IL-1 previously^31,33^, as was observed in the present study where NGF was only increased in NP cells in 3D alginate culture following addition of IL-1β to the degenerate media. It is possible that NGF and other growth factors were sequestered in the ECM and thus not fully represented by the conditioned media. Future studies should include protein quantification within digested tissue explants.

Interestingly greater effects were observed in the secretome from alginate cultures as opposed to tissue explants which could also be due to an increased response of isolated cells expanded in high glucose media and subjected to 2-week re-differentiation period prior to stimulation with differential media for 72hrs. However, it could also indicate a dampening of the effects of physiological media within tissue cultures as opposed to 3D alginate cultures. This could be due to the fact that the cells reside within an intact pericellular matrix in NP tissue explants, whilst in alginate only limited pericellular matrix is deposited over the re-differentiation period^26^, insufficient to match that of native tissue, a phenomenon also observed in isolated chondrocytes v/s cartilage tissue cultures^57^. Alternatively the reduced response in tissue explants could be a result of reduced nutrient and waste diffusion across tissue explants, particularly as the cultures within the current study were not biomechanically loaded^58^.

At the matrisome level, it was notable that even within alginate cultures at a concentration of 100pg/ml, IL-1β failed to induce an increase in the matrix degrading enzymes investigated. This lack of stimulation was also seen within the degenerate NP explant cultures in the current study and previously^39,42^. Similarly, within our recently published study investigating the potential of IL-1R antagonist (IL-1Ra) gene therapy where human NP explants were cultured within low glucose media under physioxia (equivalent to the standard media condition in the current study), supplementation with 100pg/ml IL-1β did not induce matrix degrading enzymes, although inhibition of IL-1 via IL-1Ra inhibited catabolism^39^. Concentrations of IL-1β identified in the secretome of human tissue explants in our previous study ranged from 108–526pg/mL, median: 253pg/mL IL-1β^39^, and thus addition of 100pg/ml IL-1β would not substantially increase the overall concentration of IL-1β, whilst concentrations of IL-1β identified within standard culture conditions of 3D alginate beads within the current study ranged between 0.3-2pg/ml and thus upon addition of 100pg/ml IL-1β to these cultures represented an increase in IL-1β concentrations to those predicted to be seen *in vivo.* Within these alginate bead cultures the addition of IL-1β resulted in a further stimulation of IL-1β, with concentrations of IL-1β within conditioned media increasing to 200-300pg/ml, indicating an increased production of IL-1β following stimulation with IL-1β. Such a positive feedback loop has previously been shown in NP cells isolated from degenerate but not healthy NP tissues when stimulated with IL-1β^29^.

### Study Limitations

A key element which remains missing in the current study is the application of physiological loading. Which can be applied using bioreactor systems designed to load tissue explants^59,60^ or whole organ systems^61^. Thus, it would be recommended to apply the physiological media developed here together with the sophisticated loading culture systems developed by groups worldwide^61^. The desired external conditions to mimic the NP condition particularly within organ cultures would need to be further optimised considering the diffusion rates for O_2_ and solutes under diurnal loading regimes applied to develop a complex model of the IVD. Furthermore, additional donor samples would be required to unpick differential responses between donor demographics and influence of degenerative state of tissues. This study aimed to develop media which mimicked more closely the environmental conditions of the native disc, however to determine the specific influences of pH, Glucose, O_2_, culture system (3D alginate, or tissue explants) and catabolic cytokines, a multifactorial design would be required to pin point potential influencers, however given the limited responses within modified media, this suggests that native disc cells are adapted to these conditions *in vivo*. This study focused on the utilisation of physiological levels of IL-1β to induce catabolic response, however limited response was observed within tissue explants, whereby cytokine levels were already high, suggesting higher doses may be required to further stimulate an increase in degenerative state. Further investigation of alternative cytokines may provide greater catabolic responses such as IL-6, TNF although these factors are also induced by IL-1.

## Conclusions

Here, the development of media compositions to mimic the healthy and degenerate IVD have been achieved. Human NP cells and explants were successfully cultured within these media without any detectable loss of viability and minimal secretomic changes. The main effects observed within cultures were seen with the addition of physiological concentrations of IL-1β within 3D cultures of expanded cells. These conditions provide appropriate environmental conditions *in vitro* which mimic more closely the intra-discal conditions observed during healthy and degenerate physiology and further studies are warranted to determine behaviour across broader patient samples. The application of these media can provide more appropriate culture conditions to test potential therapeutic approaches and understand more fully the pathogenesis of disease using *in vitro* and *ex vivo* models.

## Acknowledgments

Our sincere thanks go to the surgeons at Northern General Hospital and Claremont Hospital (Mr Ashley Cole, Mr Neil Chiverton, Mr Lee Breakwell, Mr Michael Athanassacopoulos, Mr Marcel Ivanov, Mr James Tomlinson, Miss Shreya Srinivas, Mr Surya Gandham and Mr Paul Brewer) for their invaluable efforts in collecting tissue samples, as well as to the patients for their donations. This work was supported by the funding received from the European Union’s Horizon 2020 research and innovation programme [grant agreement number 825925] (MAT, CLM), the Marie Skłodowska Curie International Training Network (ITN) “disc4all” grant agreement #955735 (CLM, LA), and the NC-CHOICE project [file number 19251] the Dutch Research Council (NWO) Talent programme VICI TTW (MAT).

## Author contributions

CLM, JS, SB and MAT contributed to conception and design of the study; JS, SB, EK and AB contributed to acquisition of laboratory data; JS, EK and CLM performed the data analysis; JS, SB, EK, LP, MAT and CLM contributed to interpretation of the data; JS, SB and CLM drafted the manuscript; All authors critically revised the manuscript for intellectual content; All authors approve the final version and agree to be accountable for all aspects of the work.

---

**Supplementary Figure 1:**
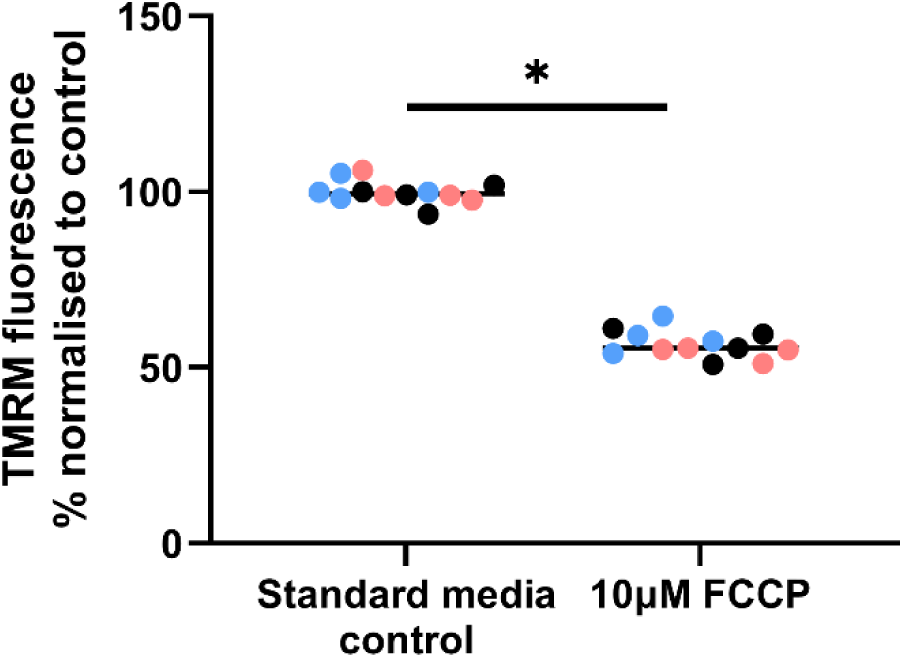
Mitochondrial activity – control vs FCCP. Mitochondrial activity determined using Image-iT™ TMRM Reagent in alginate bead-encapsulated human NP cells in standard media cultured in 5% O_2,_ 37°C, 5% CO_2_ +/− 10µM carbonyl cyanide-4-(trifluoromethoxy) phenylhydrazone (FCCP) treatment for 10 mins prior to reading TMRM fluorescence. Individual patient data points shown in differential colours. *=*p*≤0.05.

**Supplementary Figure 2:**
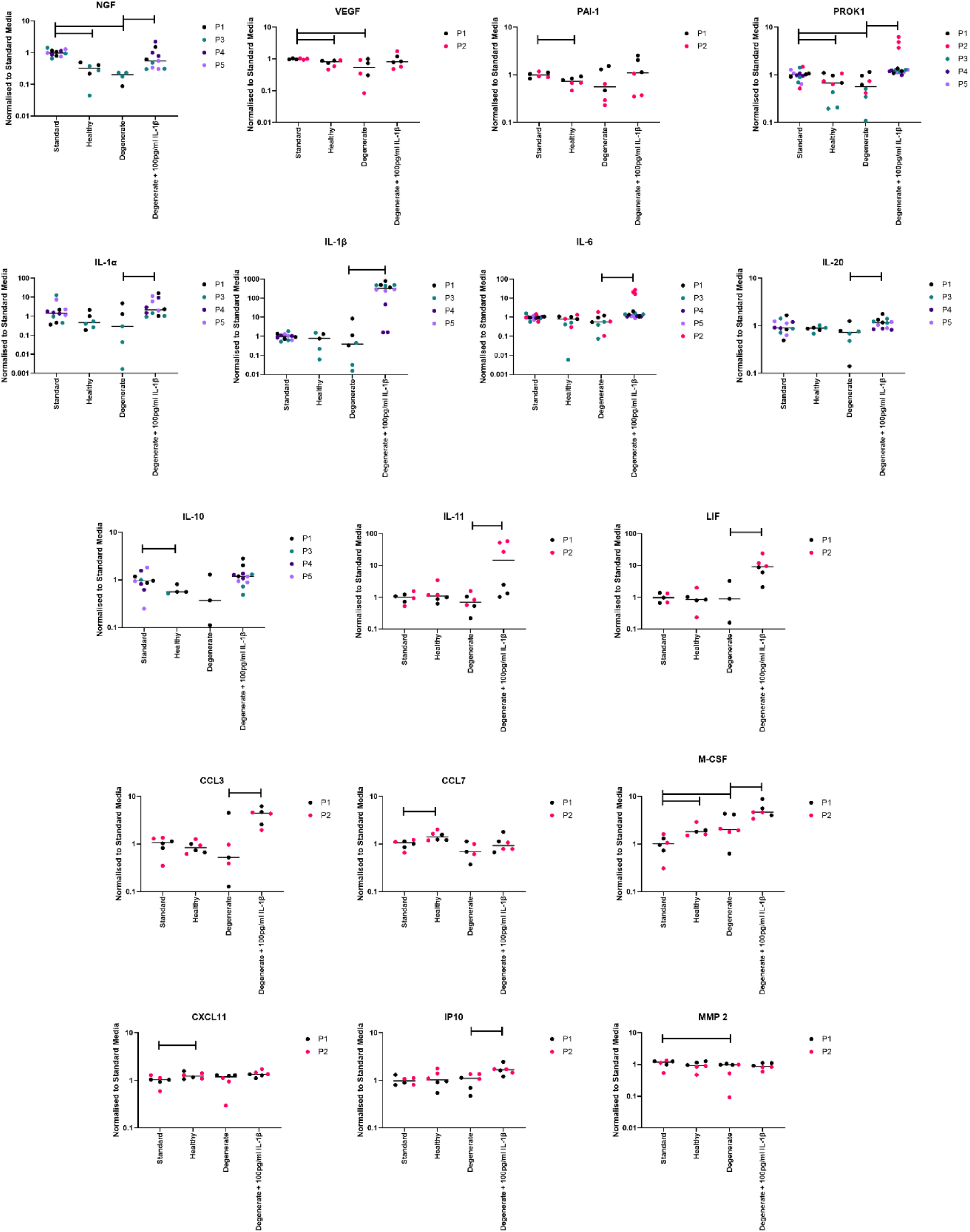
Secretome release from 3D cultured NP cells. Scatter plots shown for differentially produced proteins between culture media. Angiogenic and neurogenic proteins (NGF, VEGF, PAI-1, PROK1), Inflammatory cytokines (IL-1α, IL-1β, IL-6, IL-20) and anti inflammatory factors (IL-10, IL-11, LIF), Chemokines (CCL3, CCL7, M-CSF, CXCL11, IP10) and matrix degrading enzyme (MMP 2). *=*p*≤0.05.

## References

1. Urban JPG. 2002. The role of the physicochemical environment in determining disc cell behaviour. Biochem. Soc. Trans. 30(6):858–863.

2. Crump KB, Alminnawi A, Bermudez-Lekerika P, et al. 2023. Cartilaginous endplates: A comprehensive review on a neglected structure in intervertebral disc research. JOR SPINE.

3. Wong J, Sampson SL, Bell-Briones H, et al. 2019. Nutrient supply and nucleus pulposus cell function: effects of the transport properties of the cartilage endplate and potential implications for intradiscal biologic therapy. Osteoarthr. Cartil. 27(6):956–964.

4. Urban JP, Holm S, Maroudas A, Nachemson A. 1977. Nutrition of the intervertebral disk. An in vivo study of solute transport. Clin. Orthop. Relat. Res. (129):101–14.

5. Bibby SRS, Jones DA, Ripley RM, Urban JPG. 2005. Metabolism of the Intervertebral Disc: Effects of Low Levels of Oxygen, Glucose, and pH on Rates of Energy Metabolism of Bovine Nucleus Pulposus Cells. Spine (Phila. Pa. 1976). 30(5):487–496.

6. Urban JPG, Holm S, Maroudas A. 1978. Diffusion of small solutes into the intervertebral disc: as in vitro study. Biorheology 15(3–4):203–223.

7. Diamant B, Karlsson J, Nachemson A. 1968. Correlation between lactate levels and pH in discs of patients with lumbar rhizopathies. Experientia 24(12):1195–1196.

8. McDonnell EE, Buckley CT. 2022. Consolidating and re-evaluating the human disc nutrient microenvironment. JOR SPINE 5(1).

9. Nachemson A. 1969. Intradiscal Measurements of pH in Patients with Lumbar Rhizopathies. Acta Orthop. Scand. 40(1):23–42.

10. Adams MA, Roughley PJ. 2006. What is Intervertebral Disc Degeneration, and What Causes It? Spine (Phila. Pa. 1976). 31(18):2151–2161.

11. Johnson ZI, Shapiro IM, Risbud M V. 2014. Extracellular osmolarity regulates matrix homeostasis in the intervertebral disc and articular cartilage: Evolving role of TonEBP. Matrix Biol. 40:10–16.

12. Bermudez-Lekerika P, Crump KB, Tseranidou S, et al. 2022. Immuno-Modulatory Effects of Intervertebral Disc Cells. Front. Cell Dev. Biol. 10.

13. Roughley PJ, Mort JS. 2014. The role of aggrecan in normal and osteoarthritic cartilage. J. Exp. Orthop. 1(1):8.

14. Silagi ES, Shapiro IM, Risbud M V. 2018. Glycosaminoglycan synthesis in the nucleus pulposus: Dysregulation and the pathogenesis of disc degeneration. Matrix Biol. 71-72:368–379.

15. Snuggs JW, Bunning RAD, Le Maitre CL. 2021. Osmotic adaptation of nucleus pulposus cells: The role of aquaporin 1, aquaporin 4 and transient receptor potential vanilloid 4. Eur. Cells Mater. 41.

16. Mwale F, Ciobanu I, Giannitsios D, et al. 2011. Effect of Oxygen Levels on Proteoglycan Synthesis by Intervertebral Disc Cells. Spine (Phila. Pa. 1976). 36(2):E131–E138.

17. Naqvi SM, Buckley CT. 2015. Extracellular matrix production by nucleus pulposus and bone marrow stem cells in response to altered oxygen and glucose microenvironments. J. Anat. 227(6):757–766.

18. Razaq S, Wilkins RJ, Urban JPG. 2003. The effect of extracellular pH on matrix turnover by cells of the bovine nucleus pulposus. Eur. Spine J. 12(4):341–9.

19. Fu J, Yu W, Jiang D. 2018. Acidic pH promotes nucleus pulposus cell senescence through activating the p38 MAPK pathway. Biosci. Rep. 38(6).

20. Han B, Wang H, Li H, et al. 2014. Nucleus Pulposus Mesenchymal Stem Cells in Acidic Conditions Mimicking Degenerative Intervertebral Discs Give Better Performance than Adipose Tissue-Derived Mesenchymal Stem Cells. Cells Tissues Organs 199(5–6):342–352.

21. Hodson NW, Patel S, Richardson SM, et al. 2018. Degenerate intervertebral disc-like pH induces a catabolic mechanoresponse in human nucleus pulposus cells. JOR SPINE 1(1).

22. Arnon R. 1970. [14] Papain. Methods Enzymol. 19(C):226–244.

23. Borrelli C, Buckley CT. 2020. Synergistic Effects of Acidic pH and Pro-Inflammatory Cytokines IL-1β and TNF-α for Cell-Based Intervertebral Disc Regeneration. Appl. Sci. 10(24):9009.

24. Smith LJ, Silverman L, Sakai D, et al. 2018. Advancing cell therapies for intervertebral disc regeneration from the lab to the clinic: Recommendations of the ORS spine section. JOR Spine 1(4).

25. Phillips KLE, Chiverton N, Michael ALR, et al. 2013. The cytokine and chemokine expression profile of nucleus pulposus cells: Implications for degeneration and regeneration of the intervertebral disc. Arthritis Res. Ther. 15(6).

26. Le Maitre CL, Freemont AJ, Hoyland JA. 2005. The role of interleukin-1 in the pathogenesis of human intervertebral disc degeneration. Arthritis Res. & Ther. 7(4).

27. Le Maitre CL, Richardson SMA, Baird P, et al. 2005. Expression of receptors for putative anabolic growth factors in human intervertebral disc: Implications for repair and regeneration of the disc. J. Pathol. 207(4).

28. Johnson Z, Schoepflin Z, Choi H, et al. 2015. Disc in flames: Roles of TNF-α and IL-1β in intervertebral disc degeneration. Eur. Cells Mater. 30:104–117.

29. Le Maitre C, Freemont AJ, Hoyland J. 2005. The role of interleukin-1 in the pathogenesis of human Intervertebral disc degeneration. Arthritis Res. Ther. 7(4):R732.

30. Phillips KLE, Cullen K, Chiverton N, et al. 2015. Potential roles of cytokines and chemokines in human intervertebral disc degeneration: Interleukin-1 is a master regulator of catabolic processes. Osteoarthr. Cartil. 23(7).

31. LA Binch A, Cole AA, Breakwell LM, et al. 2014. Expression and regulation of neurotrophic and angiogenic factors during human intervertebral disc degeneration. Arthritis Res. Ther. 16(5).

32. Lee JM, Song JY, Baek M, et al. 2011. Interleukin-1β induces angiogenesis and innervation in human intervertebral disc degeneration. J. Orthop. Res. 29(2):265–269.

33. Purmessur D, Freemont AJ, Hoyland JA. 2008. Expression and regulation of neurotrophins in the nondegenerate and degenerate human intervertebral disc. Arthritis Res. Ther. 10(4):R99.

34. Binch ALA, Cole AA, Breakwell LM, et al. 2015. Class 3 semaphorins expression and association with innervation and angiogenesis within the degenerate human intervertebral disc. Oncotarget 6(21).

35. Lama P, Le Maitre CL, Harding IJ, et al. 2018. Nerves and blood vessels in degenerated intervertebral discs are confined to physically disrupted tissue. J. Anat. 233(1).

36. Binch ALA, Cole AA, Breakwell LM, et al. 2015. Nerves are more abundant than blood vessels in the degenerate human intervertebral disc. Arthritis Res. Ther. 17(1).

37. Freemont A, Peacock T, Goupille P, et al. 1997. Nerve ingrowth into diseased intervertebral disc in chronic back pain. Lancet 350(9072):178–181.

38. Basatvat S, Bach FC, Barcellona MN, et al. 2023. Harmonization and standardization of nucleus pulposus cell extraction and culture methods. JOR Spine 6(1).

39. Snuggs JW, Senter RK, Whitt JP, et al. 2025. PCRX-201, a novel IL-1Ra gene therapy treatment approach for low back pain resulting from intervertebral disc degeneration. Gene Ther. 32(2):93–105.

40. Montrose MH, Knoblauch C, Murer H. 1988. Separate control of regulatory volume increase and Na+-H+ exchange by cultured renal cells. Am. J. Physiol. Physiol. 255(1):C76–C85.

41. Le Maitre CL, Hoyland JA, Freemont AJ. 2004. Studies of human intervertebral disc cell function in a constrained in vitro tissue culture system. Spine (Phila. Pa. 1976). 29(11).

42. van Maanen JC, Bach FC, Snuggs JW, et al. 2025. Explorative Study of Modulatory Effects of Notochordal Cell-Derived Extracellular Vesicles on the IL −1β-Induced Catabolic Cascade in Nucleus Pulposus Cell Pellets and Explants. JOR SPINE 8(1).

43. Rampersad SN. 2012. Multiple Applications of Alamar Blue as an Indicator of Metabolic Function and Cellular Health in Cell Viability Bioassays. Sensors 12(9):12347–12360.

44. Le Maitre CL, Freemont AJ, Hoyland JA. 2009. Expression of cartilage-derived morphogenetic protein in human intervertebral discs and its effect on matrix synthesis in degenerate human nucleus pulposus cells. Arthritis Res. Ther. 11(5).

45. Desai S, Grefte S, van de Westerlo E, et al. 2024. Performance of TMRM and Mitotrackers in mitochondrial morphofunctional analysis of primary human skin fibroblasts. Biochim. Biophys. Acta - Bioenerg. 1865(2):149027.

46. Bermudez-Lekerika P, Tseranidou S, Kanelis E, et al. 2025. Ex Vivo and In Vitro Proteomic Approach to Elucidate the Relevance of IL −4 and IL −10 in Intervertebral Disc Pathophysiology. JOR SPINE 8(1).

47. Binch A, Snuggs J, Le Maitre CL. 2020. Immunohistochemical analysis of protein expression in formalin fixed paraffin embedded human intervertebral disc tissues. JOR Spine 3(3).

48. Cherif H, Li L, Snuggs J, et al. 2023. Injectable hydrogel induces regeneration of naturally degenerate human intervertebral discs in a loaded organ culture model. Acta Biomater..

49. Gruber HE, Watts JA, Riley FE, et al. 2013. Mitochondrial bioenergetics, mass, and morphology are altered in cells of the degenerating human annulus. J. Orthop. Res. 31(8):1270–1275.

50. Heart E, Yaney GC, Corkey RF, et al. 2007. Ca2+, NAD(P)H and membrane potential changes in pancreatic β-cells by methyl succinate: comparison with glucose. Biochem. J. 403(1):197–205.

51. Chan KK, Lee AC, Chung SR, et al. 2025. Upregulations of SNAT2 and GLS-1 Are Key Osmoregulatory Responses of Human Corneal Epithelial Cells to Hyperosmotic Stress. J. Proteome Res. 24(6):2771–2782.

52. Tök L, Nazıroğlu M, Uğuz AC, Tök Ö. 2014. Elevated hydrostatic pressures induce apoptosis and oxidative stress through mitochondrial membrane depolarization in PC12 neuronal cells: A cell culture model of glaucoma. J. Recept. Signal Transduct. 34(5):410–416.

53. Malas KM, Lambert DS, Heisner JS, et al. 2022. Time and charge/pH-dependent activation of K+ channel-mediated K+ influx and K+/H+ exchange in guinea pig heart isolated mitochondria; role in bioenergetic stability. Biochim. Biophys. Acta - Bioenerg. 1863(8):148908.

54. Song Y, Wang Y, Zhang Y, et al. 2017. Advanced glycation end products regulate anabolic and catabolic activities *via* NLRP3-inflammasome activation in human nucleus pulposus cells. J. Cell. Mol. Med. 21(7):1373–1387.

55. Zhang Z, Wu O, Ying J, et al. 2025. Regulation of diabetic disc degeneration: The role of AGEAT/miR-204-5p/Mapk4 axis in nucleus pulposus cells’ mitochondrial function and apoptosis. Cell. Signal. 133:111857.

56. Nasto LA, Robinson AR, Ngo K, et al. 2013. Mitochondrial-derived reactive oxygen species (ROS) play a causal role in aging-related intervertebral disc degeneration. J. Orthop. Res. 31(7):1150–1157.

57. Graff RD, Kelley SS, Lee GM. 2003. Role of pericellular matrix in development of a mechanically functional neocartilage. Biotechnol. Bioeng. 82(4):457–464.

58. Salzer E, Gorgin Karaji Z, van Doeselaar M, et al. 2025. The role of loading-induced convection versus diffusion on the transport of small molecules into the intervertebral disc. Eur. Spine J. 34(1):326–337.

59. Salzer E, Mouser VHM, Bulsink JA, et al. 2023. Dynamic loading leads to increased metabolic activity and spatial redistribution of viable cell density in nucleus pulposus tissue. JOR spine 6(1):e1240.

60. Salzer E, Schmitz TC, Mouser VHM, et al. 2023. EX VIVO INTERVERTEBRAL DISC CULTURES: DEGENERATION-INDUCTION METHODS AND THEIR IMPLICATIONS FOR CLINICAL TRANSLATION. Eur. Cells Mater. 45.

61. Tang SN, Bonilla AF, Chahine NO, et al. 2022. Controversies in spine research: Organ culture versus in vivo models for studies of the intervertebral disc. JOR Spine 5(4).

